# Evolutionary stability of plant-pollinator networks: efficient communities, hysteresis, and a pollination dilemma

**DOI:** 10.1101/286294

**Authors:** Soeren Metelmann, Shoko Sakai, Michio Kondoh, Arndt Telschow

## Abstract

Mutualistic interactions between species such as pollination and plant–mycorrhiza interactions are ubiquitous in nature and essential for ecosystem functioning. Often dozens or even hundreds of species with different degree of specialization form complex networks of interdependence. How the complexity evolved and is maintained are fundamental questions in ecology. Here, we present a new game theoretical approach to model complex mutualistic interactions, which we apply to pollination networks. The theoretical analysis revealed multiple evolutionary stable network structures that form a gradient from generalism toward specialism with increasing availability of pollination service. In particular, we found that efficient communities evolve only under pollination oversupply, but that pollination shortage selects for inefficient network structures due to a pollination dilemma. These results suggest that availability of pollination services is a key factor structuring pollination networks and offer a new explanation for the geographical differences in pollination faunas that have long been recognized by ecologists. The study bridges the gap between network studies, game theory, and the natural history of pollination, which have hitherto been studied largely independently.

## Introduction

Mutualistic networks between organisms from different kingdoms or domains are common in nature: Plants interact with rhizobial bacteria or mycorrhizal fungi for enhanced access to nitrogen or phosphorus, in exchange for carbon in form of photosynthesis products (Sprent & Raven 1985; Rayner 1927). The human gut flora provides us with protection against pathogenic microorganisms, as well as metabolites we would not be able to digest on our own while we supply it with nutrients (Backhed et al. 2005). Plants form nutritious fruits for birds or mammals, so that they disperse the plants’ seeds away from competing relatives and potential parasites (Howe & Smallwood, 1982). Another well-studied mutualism between plants and birds or insects are pollination networks.

For a long time, research focussed on the exclusive interactions between single plant and pollinator species (e.g. Kritsky 1991, Pellmyr & Huth 1994). However, the majority of the plant species are pollinated by a wide range of animals, while these pollinators often visit a wide range of plant species in return (Bascompte et al. 2003; Ollerton et al. 1996). Resulting pollination networks can span hundreds of plant and animal species (Herrera 1988; Primack 1983). An important aspect of such a network is that its structure strongly affects the fitness of individual plants (Muchhala et al. 2010). Pollinators that are specialized to a certain plant species have a high rate of conspecific pollen transport, whereas the efficiency of pollination is low for generalist pollinators that are shared by different plant species (Bell et al. 2005; Wheelwright & Orian 1982). While the cost of sharing pollinators has been recognized as an important factor for plant specialization (Muchhala et al. 2010; Sargant & Otto 2006), it has rarely been explicitly connected to the network structure itself (Mitchell et al. 2009).

Since fitness of a plant depends on not only the strategy of the species but also strategies of other plant species in the same community, the situation can be regarded as a game between plant species. Following previous work on co-evolutionarily stable strategies and networks (Matsuda and Namba 1991; Roughgarden 1969), we designed a game theoretical model in which the players are the different plant species in the community. We use plant fitness as payoff, measured as the efficiency of conspecific pollen transport. The theoretical analysis revealed multiple evolutionary stable network structures that differ in the level of pollination efficiency and form a gradient from generalism toward specialism with increasing availability of pollination service. These results suggest that availability of pollination services is a key factor structuring pollination networks.

## Material and methods

### Game theoretical model

A community consisting of *N* plant and *M* animal species is described by a symmetric game in which the plants species are the players. They can form or abort links to pollinators and the strategy space for each plant species consists of all combinations of links to the *M* animal species. The N plant and M pollinator species are assumed to have constant population sizes, denoted by *p*_*i*_ and *a*_*j*_ for plant species *i* and animal species *j*, respectively. Once the network structure is determined by the *N* plant strategies, each animal individual visits two plants that are randomly chosen from all plant individuals to which the animal is linked. Pollen uptake happens at the first visit and pollination at the second plant.

Each plant individual is assumed to produce a certain amount of pollen (*r*), which is removed by its visitors with equal probabilities. However, because of limited uptake by each animal individual (*u*), some pollen may be left on the plants. The payoff for each plant species (*Q*_*i*_) is then calculated as the average fraction of its pollen that is transported conspecifically during pollination. *Q*_*i*_ is a measure based on the male component of plant fitness and describes the efficiency of pollination of plant species *i*.

For a given network structure, the pollination efficiency *Q*_*i*_ is calculated as follows. Let ***L*** be a *N*×*M* matrix whose elements are defined as *L*_*i,j*_ = 1 if plant species *i* is pollinated by animal species *j*, and *L*_*i,j*_ = 0 otherwise. Then, the total number of animal individuals visiting plant species *i* for pollen uptake computes to 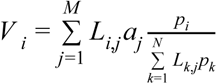 Here, 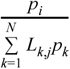 is the probability that an animal individual of species *j* visits plant species *i*, and the sum is taken over all animal species pollinating plant species *i* weighted by the animal population sizes.

The *V*_*i*_ animal visitors remove either all or only a part of the pollen of plant *i*. Note that the total pollen production by this plant species is *rp*_*i*_. As each animal individual has a limited amount of pollen uptake *u*, the maximal possible pollen uptake from plant species *i* is *uV*_*i*_. Accordingly, there is a pollinator shortage for *uV*_*i*_<*rp*_*i*_ and a pollinator oversupply for *uV*_*i*_>*rp*_*i*_. These two cases must be distinguished to calculate the pollination efficiency *Q*_*i*_. For a pollinator shortage, it holds that

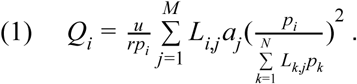

The sum on the right-hand side of equation (1) is the total number of animals that visit plant species *i* twice, once for pollen uptake and once for pollen delivery. Multiplication by *u* yields the total amount of pollen that is delivered conspecifically, and further division by *rp*_*i*_ gives the pollination efficiency *Q*_*i*_. For pollinator oversupply, the pollination efficiency computes to

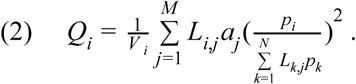

The sum on the right-hand side is the same as in equation (1). Differences occur with respect to the total amount of conspecifically-delivered pollen, which is calculated by multiplying the sum in equation (2) by *rp*_*i*_*/V*_*i*_. The factor *rp*_*i*_*/V*_*i*_ is interpreted as the average amount of pollen taken up by the animal visitors under pollination oversupply. As above, division by *rp*_*i*_yields the pollination efficiency *Q*_*i*_.

### Game theoretical analysis

The game with two plant and two animal species was analyzed by analytically calculating payoff matrices as function of animal species abundance (*a*=*a*_1_=*a*_2_). The payoff matrices were then used to determine evolutionarily stable strategies (ESS).

The plant pollinator game could not be analyzed analytically for arbitrary numbers of animal and plant species, as the number of possible pollination networks increases exponentially with N and M (2^N·M^). Therefore, an analysis for high species numbers was conducted numerically using a discrete time model that simulated the evolution from unstable to stable networks. The model was implemented in *Python 2*.*7*. A second program was written independently in *Perl 5*.*16*.*2* to test the accuracy of the results. Each run started with a community in which plant and animal species were linked randomly, and during each run all parameters were kept constant. In each time step, one plant species was randomly chosen and a link was randomly attached to or removed from it. The change was accepted if the pollination efficiency (Q_i_) of the chosen plant species increased and rejected if it decreased. If pollination efficiency stayed the same the change was accepted with a chance of 50%. This procedure was repeated 10,000 times to reach equilibrium. Each parameter constellation was repeated 100 times to avoid stochastic artefacts. We kept r=1 and u=1 for the analysis and simulations.

## Results

First, we analytically examined the plant–pollinator game with two plant and two animal species (Fig. 1). There were two stable states, and the stability depended on the abundance of pollinators in the community. Each of these stable states was associated with a certain network structure, the so-called evolutionarily stable network structure (ESNS). The analysis revealed that the ESNSs formed a gradient from generalism (fully connected network, (G, G) in Fig. 1b) towards specialism (one-to-one relationships, (S, S) in Fig. 1b) with increasing pollinator abundance.

**Figure 1.**
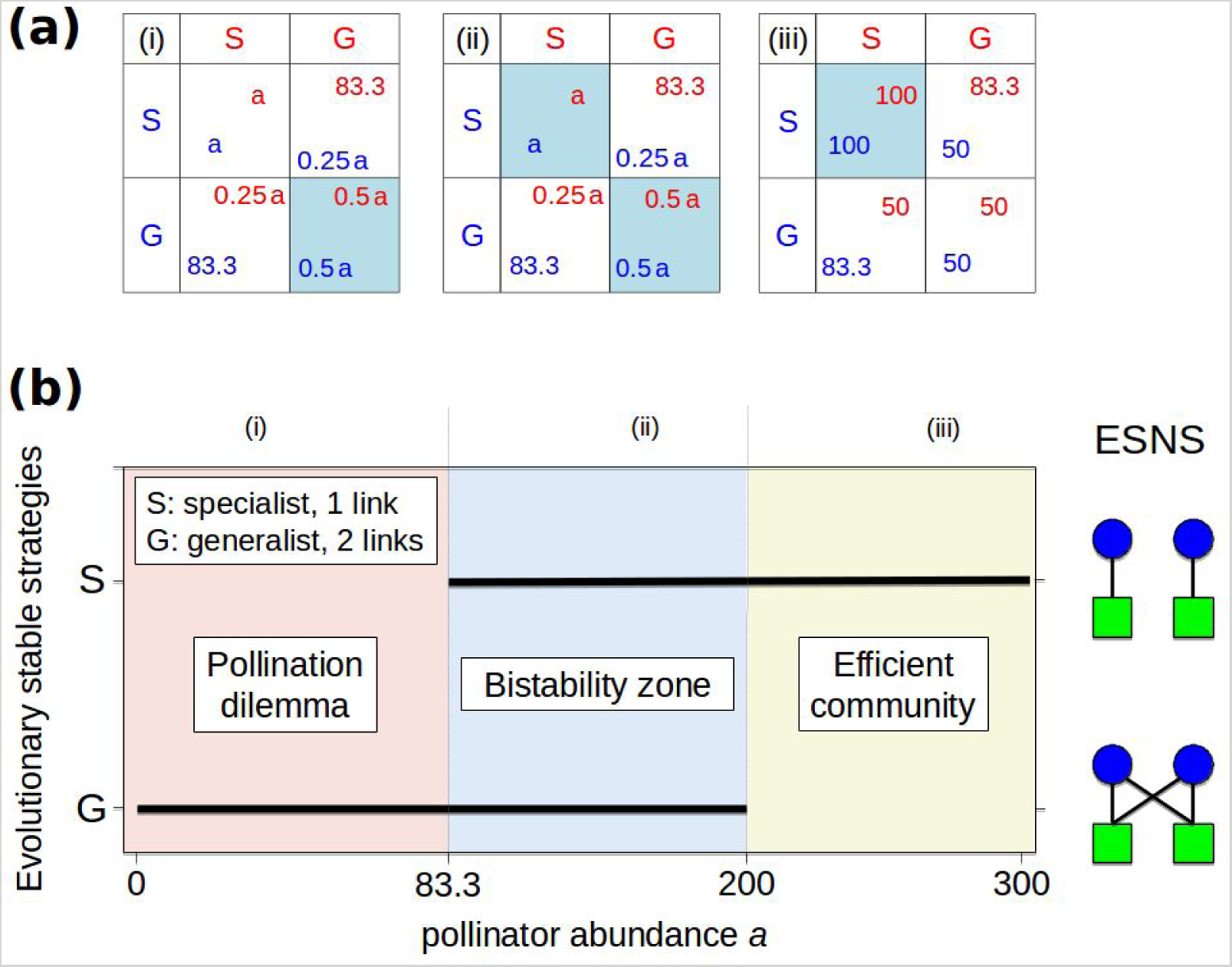
Pollination game for two plant and two animal species. There are two possible plant strategies: (S) specialist, with one link; (G) generalist, with two links. The payoff is measured as percentage of produced pollen that is transferred conspecifically. **(a)** Payoff matrices as a function of pollinator abundance *a* (*a*=*a*_1_=*a*_2_; *a*_*i*_, abundance of pollinator species *i*). The payoff of plant species 1 and 2 are indicated by blue and red colors, and the colored boxes indicate evolutionarily stable strategies (ESS). There are three cases: (i) 66.7≤*a*<83.3; qualitatively similar payoff matrices for 0<*a*<66.7; (ii) 88.3<*a*<200; (iii) a>200. Parameters: pollen production rate per plant, *r*=1; pollen uptake rate per pollinator, *u*=1; plant population size of species 1 and 2, *p*_1_= *p*_2_=100. **(b)** Ranges of pollinator abundance where each network structure is evolutionarily stable. The right-hand side shows the evolutionary stable network structures (ESNS).

Variation in the ESNS resulted from the interplay of two costs that affect plant fitness. These costs are incomplete pollen uptake, which occurs when pollinator visits are too few to remove all pollen, and ineffective heterospecific pollen transport, which occurs when different plant species share pollinators. When pollinators are sufficiently abundant and plants do not suffer from incomplete pollen uptake (iii in Fig. 1b), plants segregate pollinators to avoid heterospecific pollen transport, and each pollinator is connected to only a single plant species. This results in an ESNS with few links (networks of (S, S) in Fig. 1b). With lower pollinator abundance (i, ii) however, the first cost overrides the second, and each individual plant can increase the amount of dispersed pollen by attracting more animal pollinator species. The resulting ESNSs are characterized by a high number of links (network of (G, G) in Fig. 1b).

Interestingly, the payoff matrix for severe pollinator shortage (i) shares basic features of the prisoner’s dilemma game. In this case, the fitness for both plant species would be highest if both segregate, but such a network structure is not evolutionarily stable. Thus, when pollinators are scarce, plants suffer not only from pollinator shortage but also the inefficient use of pollinators due to a “pollination dilemma”. Intermediate pollinator abundance (ii) allows for two stable states, one where pollinators are efficiently used in a community ((S, S) in Fig. 1b) and one “dilemma state” ((G, G) in Fig. 1b). From a game theoretical perspective, plant specialization (S) can therefore be considered as a cooperative strategy and generalism (G) as a non-cooperative one, because the use of the former results in higher pollination efficiencies in the community than the latter.

Next, we investigated the plant–pollinator game for four animal and four plant species using numerical iterations (Fig. 2). We found six evolutionary stable network structures that form a gradient from generalism towards specialism with increasing animal abundance. As above, there are three qualitatively different zones: (1) inefficient generalist networks evolve for low animal abundance due to a pollination dilemma, (2) multiple stable network structures occur for intermediate animal abundance, and (3) efficient communities with specialist interactions evolve under animal oversupply. An interesting aspect of the pollination game with four animals and plants was the emergence of network modularity (network associated to I_1_ in Fig. 2a).

**Figure 2.**
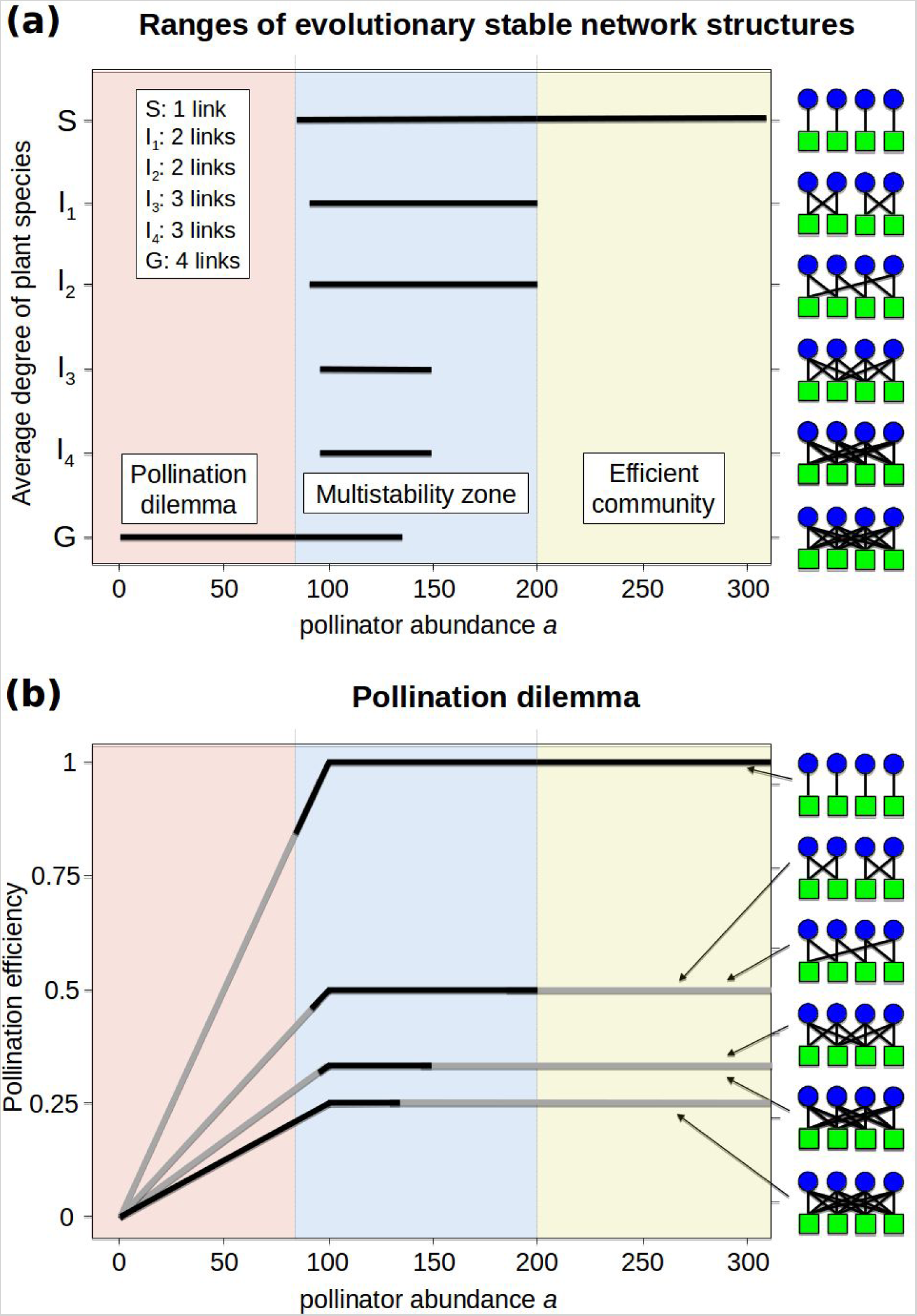
Pollination game for four plant and four animal species. **(a)** Ranges of pollinator abundance where each network structure is evolutionarily stable. The right-hand side shows the evolutionary stable network structures (ESNS) with plants as blue circles and pollinators as green squares. **(b)** Pollination efficiency as function of pollinator abundance for evolutionary stable network structures. Black and gray lines indicate parameter ranges, where the network structure is stable and unstable, respectively.

Finally, we analyzed larger networks with 20 plant species and variable animal abundance or animal species number (Fig. 3 and 4). In general, the simulation analysis revealed the same basic trends as above: (1) the evolved network structures form a gradient from generalism towards specialism with increasing animal abundance, (2) inefficient generalist networks evolve for low animal abundance due to a pollination dilemma, (3) multiple stable network structures occur for intermediate animal abundance, and (4) efficient communities with specialist interactions evolve under animal oversupply.

**Figure 3.**
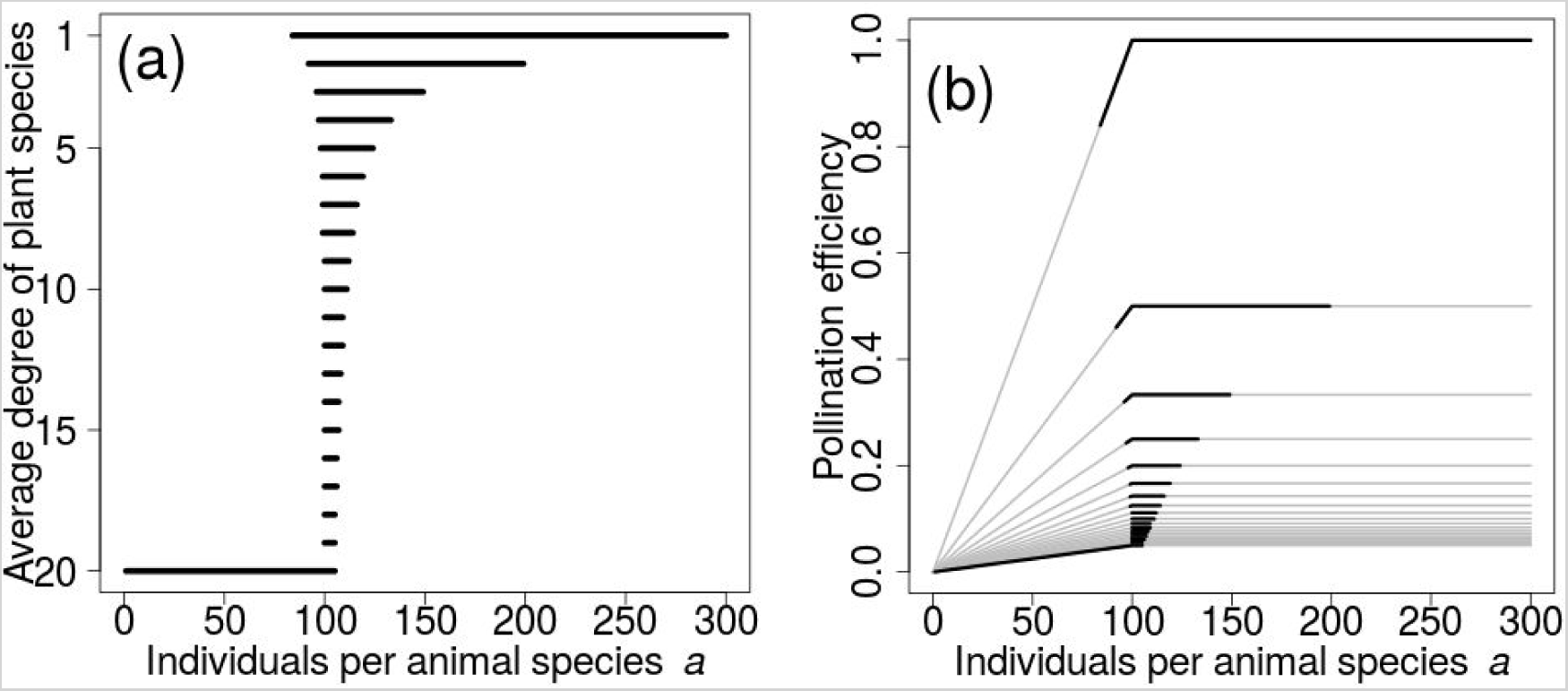
Pollination game for twenty plant and twenty animal species. **(a)** Ranges of pollinator abundance where each network structure is evolutionarily stable. **(b)** Pollination efficiency as function of pollinator abundance for evolutionary stable network structures. Black and gray lines indicate parameter ranges, where the network structure is stable and unstable, respectively.

**Figure 4.**
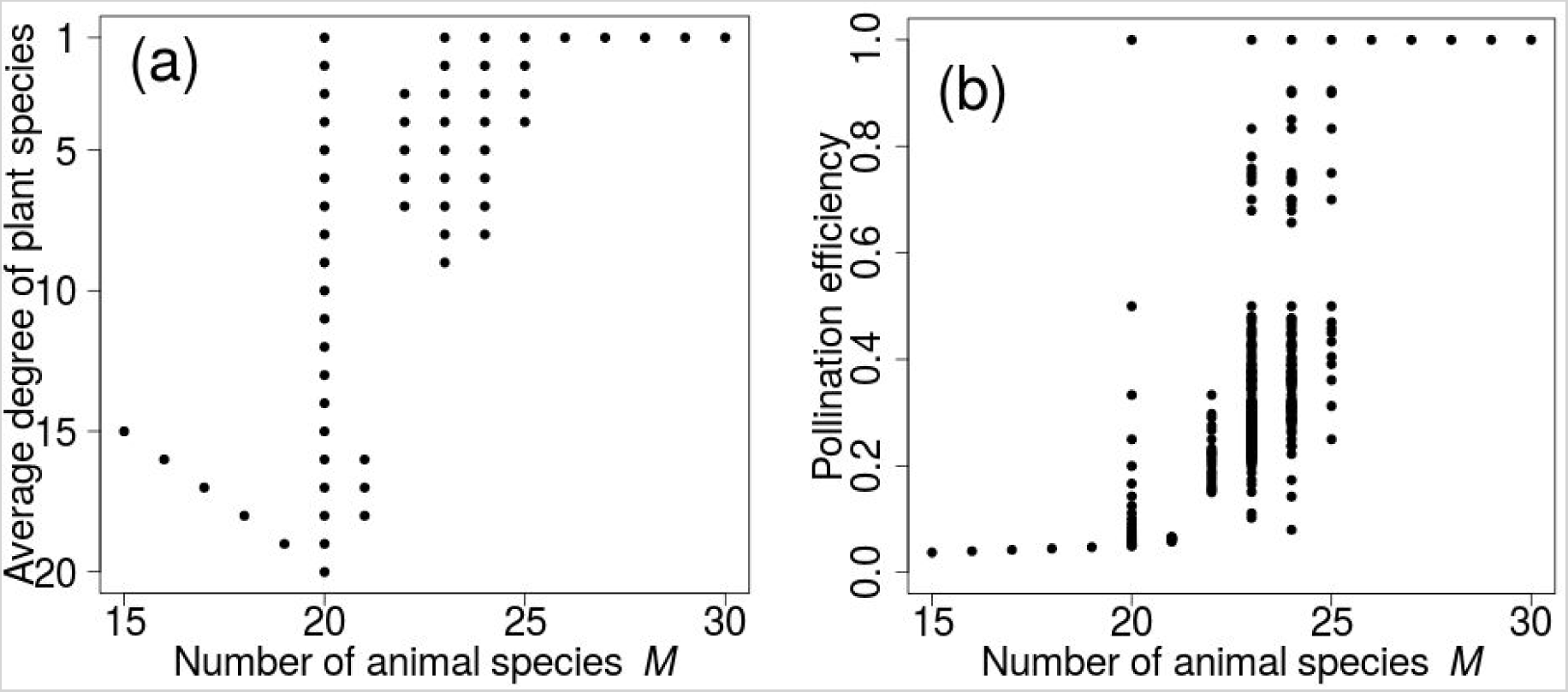
Pollination game for twenty plant species and a varying number of animal species. For each M, we tested initial networks with 1…M links per plant species. Links were distributed equally and symmetrically among pollinators, with a maximum of one link difference between them to start with “symmetrical” networks. **(a)** Number of links per evolved network structures after 50,000 steps or until structures are stable. For M > 20, we classified plants with a pollination efficiency of 1 but more than one pollinator species as having only a single pollinator species to remove artificially inflated networks **(b)** Pollination efficiency as function of pollinator abundance for evolutionary stable network structures. Both plots show irregular behavior for M=20. Here, initial networks are already stable and thus do not evolve, compare Figure 3 a).

## Discussion

### Efficient communities & pollination dilemma

To our knowledge, the structure of pollination networks have never been studied in the light of game theory. However, there are a limited number of studies that model plant specialisation as a function of pollinator abundance. One example is the one plant - two pollinator model by Waser et al. (1996). The model uses more general terms for the visiting frequency and pollination efficiency, but we can transform it into a two plant-two pollinator game with 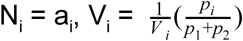 and 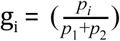, using r=1 and u=1.

In this model approach, the game was only played by the plant species and pollinators simply follow the links offered by plants. But different pollen and nectar rewards would provide good incentives for the pollinators to favor certain plant species over others (Hicks et al. 2016). However, as travel to flowers is costly and discriminating between flowers takes time (Bateman 1947; Smithson and Giroud, 2003) while on the other hand accessing different types of flowers is not costly (Waser & Ollerton 2006), pollinators should rather try to visit every flower they find and thus favor generalism.

Besides pollinator choice, other factors such as the cost for pollen and nectar production (Pélabon et al. 2012) could be taken into account for the plant’s payoff in more elaborate models. This would be interesting to follow up, as the pollination dilemma only arises from a certain combination of values in the payoff matrix (Kollock 1998). Moreover, spatial or temporal dynamics could be analysed (Waser et al. 1996). When plant species dominate in an area or season, chances are high that pollinators visit the same species consecutively, transporting its pollen conspecifically even when the pollinator is actually a generalist. Such spatial or temporal clustering would actually favor generalism in plants.

### Application for other mutualistic networks

We presented a new game theoretical approach for the analysis of pollination networks. An important question is whether other ecological networks that also experience a cost of sharing interaction partners, could be analyzed in a similar way. One example could be plant-mycorrhiza networks in which fungi can interact with several plants and vice versa (Wang & Qiu 2006), forming networks that can contain dozens of fungi or plants (Bever et al. 2001, Toju et al. 2014). It was shown that the association between fungi and plants is non-random, with fungi ranging from specialists to generalists (Husband et al. 2002), while plants tend to be generalists (Smith and Read, 2002). In mycorrhizal networks, the fungi interact with the root systems of plants in that they transfer nitrogen, phosphorus, and water to, while receiving carbon (in form of photosynthesis products) from the plant (Olsson et al. 2010). However, the degree with which plants supply their fungi with carbon varies (Zheng et al. 2014), up to the point at which they actually extract carbon from the network, supplied by the fungi or other connected plant species (Bjorkman et al. 1960; Bruns et al. 2002; Hynson et al. 2011). In this case, the according game with the plants as players would be asymmetric, as strategies (links to fungi) depend on which plant is playing.

Note that game theory approaches have been applied to plant-mycorrhizal interactions, using plant and fungi as players to analyse parasitic behaviour such as myco-heterotrophy (Steidinger and Bever, 2014).

### Geographic variation of pollination networks

Finally, we related our results to the global variations in pollination faunas. Recently, the degree of specialization in pollination networks was reported to increase from tropical toward temperate latitudes (Schleuning et al. 2012). Further, several studies have shown that generalism is more prevalent in pollination interactions on islands than on mainlands (Barrett et al. 1996; Kaiser-Bunbury et al. 2010). The model can explain both patterns under the assumption that the tropics and islands have a pollinator shortage relative to temperate regions. In terms of Figure 1, the tropics and islands would be represented by case (i) and temperate regions by cases (ii) and (iii). In other words, we hypothesize that tropics and islands suffer from a pollination dilemma, whereas pollination oversupply selects for efficient communities in temperate regions.

Although geographical variation in pollinator abundance has never been tested directly, there are some studies supporting this hypothesis. The frequency of animal-pollinated plants is higher in the tropics (Regal 1982), which could lead to pollinator shortages (Schemske et al. 2009). And Vamosi et al. (2006) could show that regions with a high species richness, which are often found in the tropics (Pianka 1966), are associated with pollinator shortage. Further, comparative studies on fruit set and species diversity suggest a more pollination-limited plant reproduction in the tropics and on islands (Larson and Barrett 2000; Tremblay et al. 2005). Finally, the higher frequency of vertebrate pollination in the tropics and on islands could be interpreted as adaptation by plants to pollinator shortages (Olesen et al. 2010; Bawa 1990; Faegri and van der Pijl 1996). Pollinator paucity might therefore be a common feature of the tropics and islands. In conclusion, the game theoretical model offers a biologically plausible and consistent explanation for the observed geographical variation in pollinator fauna.

Our results have important ramifications for (ecological) network theory. First, the approach to compare different geographical regions allows for testing hypotheses of network evolution because each community can be considered as result of a “natural experiment”. Second, the game theoretical model suggests that pollination networks are in different evolutionarily stable states depending on pollinator abundances as has been suggested previously for one or two plant species (Muchala et al 2010; Sargant and Otto 2006; Holland and Fleming 2002; Holland et al. 2005). This raises the question whether other interaction networks also have multiple evolutionarily stable states and can be analyzed by game theory. Finally, we argue for the importance of including natural history in network modeling. Pollination biologists and other ecologists have accumulated a large body of detailed information on species interactions. Incorporating this knowledge in network models is needed for developing a more profound theory of ecological interactions (Tewksbury et al. 2014).

## Acknowledgements

We thank P. Hammerstein, A. Onuma for discussions; M. Ushio and Jakob Strauß for analytical assistance; and S. Armitage and T. Kiers for comments on the earlier version of the manuscript. Funding to SK was provided by the Research Institute for Humanity and Nature Project (P5-3), FY 2011 Researcher Exchange Program between JSPS and DAAD, and the Ministry of Education, Science, Sports and Culture (no. 20405009), Japan. AT was supported by the German Science Foundation (SPP 1399, TE 976/2-1) and the Volkswagen Foundation’s evolutionary biology initiative. AT and MK were supported by the German Science foundation (TE-976/4-1).

